# A genetic model of copulatory wounding in *Drosophila melanogaster*

**DOI:** 10.64898/2026.05.29.728886

**Authors:** Sophie L. Jalkut, Lydia M. Bischoff, Maximilian J. Pitsch, Vicki P. Losick

## Abstract

In the animal kingdom, copulatory wounds occur due to aggressive behavior or injury from the female or male appendages in contact. Copulatory wounds are a source of sexual conflict as the injury can lead to death or infection. In the laboratory, *Drosophila melanogaster* does not ordinarily exhibit signs of injury during copulation, but here we find can be induced to do so by overexpression of the serine protease, *Hayan*, in either the male or female fruit fly. Melanin deposits mark sites of injury along the female ventral abdomen and occur following a single copulation event. The male fly mid-legs are the source of injury as they rub against the ventral female abdomen. Surgical removal of the male mid-legs prevents melanization while still permitting mating to occur. Likewise, removal of the right or left mid-leg results in a corresponding loss of melanization on the female abdominal side that lacks the mid-leg contact. Thus, it appears that overexpression of *Hayan* alters the female and male anatomy to sensitize it to injury during copulation. We show that these copulatory wounds heal similar to a puncture wound via the generation of enlarged multinucleated, polyploid cells. However, the size of the injury dictates the mechanism of healing. The polyploid cells arise solely by cell fusion in response to small injuries (<1,000μm^2^), whereas the endocycle is simultaneously simulated by larger wounds (>3,000μm^2^) to compensate for cell loss. Therefore, we speculate that wound-induced polyploidization may have evolved to repair similar injuries resulting from sexual conflict in nature.

## Introduction

In nature, insects regularly sustain bodily damage from predator-prey interactions, environmental factors, parasitism, and mating partners. In the case of mating, these wounds can occur within the female’s reproductive tract as seen in the seed beetle, *Callosobruchus maculatus*, or outside female genitalia in the common bed bug, *Cimex lectularius* (Stutt and Siva-Jothy, 2001, Yan et al., 2024). An injury that arises due to mating is defined as a copulatory wound (Reinhardt et al., 2015). Copulatory wounds are a source of sexual conflict as they can lead to death or infection in the mating partner that sustains the injury. Both lab reared and wild collected *Drosophila* species have been reported to exhibit signs of injury in the female reproductive tract as well as the ventral abdomen (Reinhardt et al., 2015, Subasi et al., 2025, Subasi et al., 2024). The spiny ends of the male basal genitalia can pierce the female reproductive tract during copulation; however, the cause of ventral abdominal wounds remains unknown.

In studying the role of melanization in *Drosophila melanogaster* wound healing, we unexpectedly found that overexpression of the serine protease *Hayan* in the female or male fruit fly is sufficient to induce copulatory wounding. *Hayan* is part of the Toll signal transduction pathway, which responds to microbial ligands and activates phenoloxidase, leading to the generation of melanin at the site of injury or infection (Dudzic et al., 2019, Dudzic et al., 2015, Nam et al., 2012). The overexpression of *Hayan* appears to alter the male fly mid-legs and/or the female ventral abdomen, such that upon contact, injuries arise. The copulatory wounds are akin to a needle puncture wound and heal through the formation of multinucleated, polyploid cells that are necessary to compensate for cell loss. We propose that the overexpression of *Hayan* creates a genetic model of copulatory wounding, which can be used to further elucidate the source of sexual conflict in the animal kingdom.

## Materials & Methods

### *Drosophila* husbandry and strains

The *Drosophila melanogaster* strains used in this study were raised on corn syrup, soy flour-based fly food (Archon Scientific) at 25°C with 60% humidity and 12:12 hour light and dark cycle. *Drosophila* strains were purchased from Bloomington Drosophila Stock Center as noted in the Table S1.

### Copulation Assays

Females were collected as virgins and aged 2-3 days in isolation. Virgin female flies were then combined with males and the percentage of females with melanin deposit was recorded at times indicated. For a single 1:1 mating, one virgin female fly was added to a food vial with one male fly. Copulation was visually confirmed and male and female flies were separated indefinitely after uncoupling. After ∼3 hours, pairs that had not initiated copulation were separated and copulation was considered unsuccessful. Brightfield images of females were captured daily for three days post copulation. Representative brightfield images were taken with the Olympus S2X7 stereo microscope with DP22 camera and CellSens imaging software.

### Male Leg Removal and Imaging

Adult male flies were collected, isolated and aged 2-3 days. Males were anesthetized with carbon dioxide and positioned with their ventral side up. Using spring scissors, the leg or legs of interest were cut above the tibia-femur joint. Males were given one day to recover before initiating a mating. Forelegs (with the sex comb) and mid-legs were mounted on a glass slide and imaged with Plan-NEOFLUAR Z 1x objective on the Zeiss Axio Zoom V16 microscope.

### Copulation Videos

Courtship and copulation were recorded in cylindrical courtship chambers (10 mm wide and 3 mm height), fabricated as detailed (Boutros et al., 2017). One male and one virgin female (3-4 days old) were added to a courtship chamber using a fly aspirator. Males and females were kept separate for at least 2 days prior to video capture. Videos were recorded using a Digital Coin Microscope. Copulation duration videos were collected as described (Gautham et al., 2024).

### Tissue dissection, immunostaining, and fluorescent imaging

Adult flies were collected as virgins, isolated and aged until 2-3 days old. Then male and female flies were combined and allowed to copulate for 1-2 days. Female flies were again isolated and dissected at days post copulation or injury with a stainless steel minutien insect pin (Fine Science Tools). Female flies were dissected as previously described (Bailey et al., 2020). Dissected abdomens were fixed with 4% paraformaldehyde for 30 min, permeabilized in 1× PBS with 0.3% Triton X-100 and 0.3% BSA, and then stained with primary and secondary antibodies as noted in Table S2. Immunostained abdomens were mounted in Vectashield (Vector Labs, H-1000) on a glass coverslip and slide, with the inner tissue facing out. Tissue samples were imaged on a Zeiss Axiovert with ApoTome using a 20× or a 40× dry objective. Full Z-stack images were taken at 0.5μm per slice and flattened into a SUM or MAX of stacks projections using FIJI/ImageJ (SCR_002285) imaging software for all analysis.

### Ploidy quantification

*Drosophila* ventral abdomens were immunostained to identify and measure epithelial nuclear ploidy as previously reported (Bailey et al., 2020). The epithelial nuclei in the uninjured, *epi>+* strains were used as an internal control as ploidy was previously measured to be diploid (2C). All samples were imaged under the same conditions and settings. Using FIJI/ImageJ, epithelial nuclei were identified based on Grh expression and DAPI intensity measured. The nuclear ploidy was calculated by normalizing the DAPI intensity of the average value of the 2C uninjured epithelial nuclei for at least 3 abdomens per condition. The normalized ploidy values were binned into groups: 2C (0.6 – 2.99C), 4C (3.0C -5.99C), >6C (>6.0C).

### Cell and melanin size analysis

The syncytium and melanin deposit size were quantified by outlining the area using FIJI/ImageJ software. For syncytium, the FasIII cell junction marker served as a guide. The number of epithelial nuclei within the syncytium were quantified using the cell counter tool based on the number of Grh+ nuclei within the syncytial cell border. The melanin overlay images were generated by cropping the abdomen (300 x 400 pixels), centering the image on the middle sternites, and straightening the images as needed. Regions of interest were then drawn around each melanin deposit and overlaid onto an abdominal image of a virgin fly in the same strain.

### Replicates and statistical analysis

All experiments were performed in triplicate with at least 3 biological replicates (fruit flies) per experiment. Raw values from each graph are available in the Source Data. Excel was used for basic calculations (ex. ploidy, syncytium size), and statistical analysis was performed using GraphPad Prism (i.e. ANOVA with Tukey’s multiple comparisons test and Student *t*-test with Welch’s correction) as indicated in the figure legend.

## Results & Discussion

Melanin is generated through the oxidation of phenols to quinones, which is catalyzed by phenoloxidase (PO). The serine proteases *Hayan, MP1*, and *MP2/Sp7* have been shown to cleave prophenoloxidase (PPO) into its active PO form in response to injury (Figure 1A) (Nam et al., 2012, Tang et al., 2006). Overexpression of the individual serine proteases using the Gal4/UAS system is sufficient to induce melanin deposits in larval development. However, the ubiquitous activation of PO often results in lethality (Nam et al., 2012, Tang et al., 2006). To study the effect of melanization on the adult fly epithelium, we sought to restrict its expression using a ventrally enriched epithelial Gal4 driver (GMR51F10, henceforth *epi>*) (Losick et al., 2013, Bailey et al., 2020). In doing so, we found that only the engineered expression of *Hayan* (*epi>Hayan*) caused melanin deposits in the ventral abdomen (Figure 1B). Melanin was observed in 79% of 3-day old *epi>Hayan* flies, whereas *epi>MP1* and *epi>Sp7* flies failed to develop melanin deposits. We also noticed that whereas female *epi>Hayan* flies had melanin deposits, their male counterparts did not (Figure 1C). Therefore, it appears that *Hayan* expression with the *epi>* driver results in female-specific melanization.

**Figure 1.**
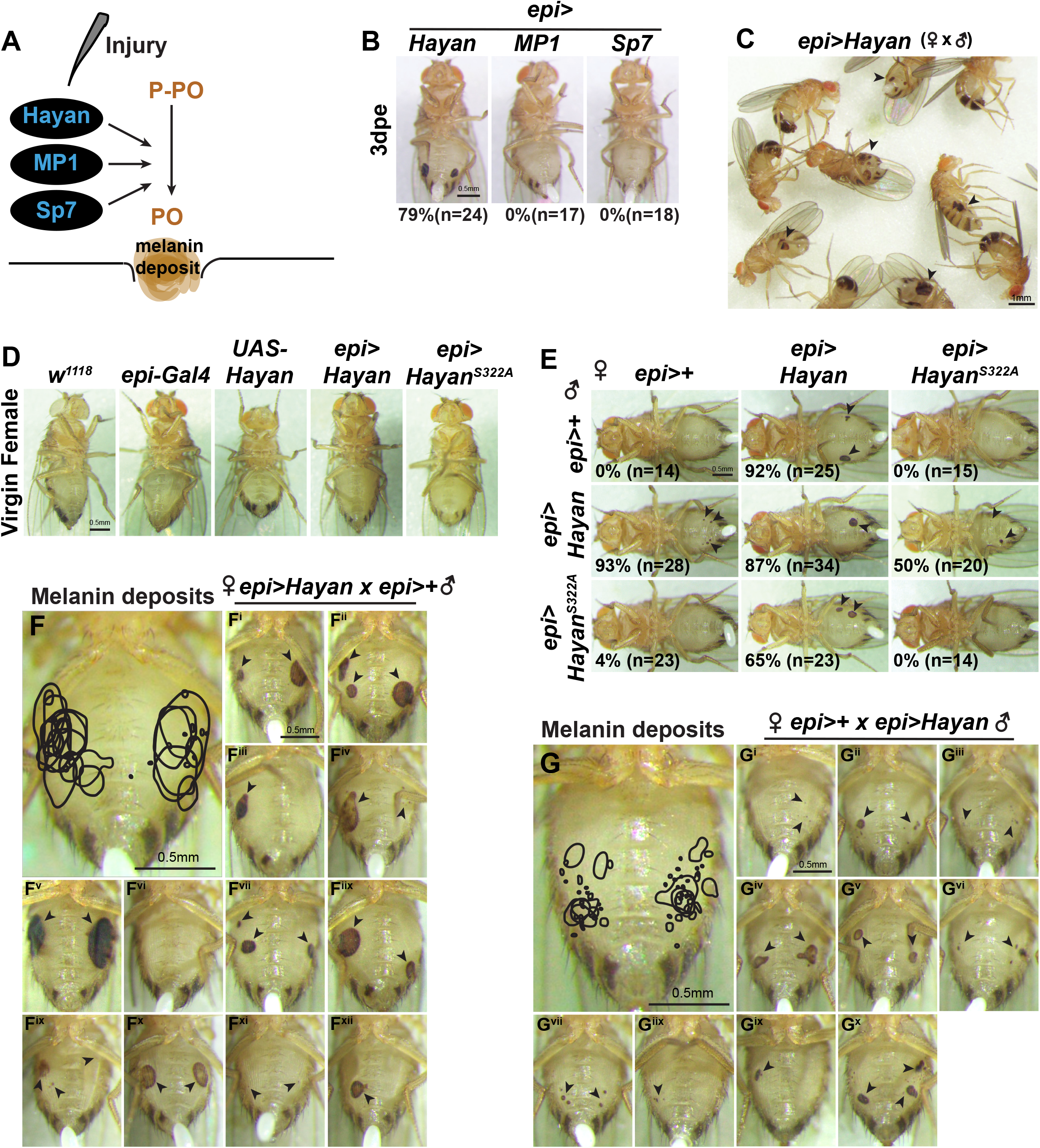
Copulation-induced melanization arises on the female ventral abdomen after a single mating when either a male or female fly overexpresses *Hayan*. (A) An illustration of the melanization pathway in *Drosophila*, where serine proteases, Hayan, MP1, or MP2/Sp7, cleave prophenoloxidase (PPO) to phenoloxidase (PO) generating melanin at the site of injury. (B) Brightfield images of female fruit flies overexpressing serine proteases via GMR51F10-Gal4 (*epi>*) at 3 days post eclosion (dpe). (C) Male and female flies expressing *Hayan*, however only the female flies exhibit melanization on their ventral abdomens (arrowheads). (D) Virgin female flies in strains indicated at 2-3dpe. (E) A 9-way mating scheme where flies were paired 1:1 and mated female flies were scored based on percent melanized at 3 days post copulation (dpc). (F and G) Melanized regions of interest (ROI) from mating pairs indicated. The ROIs were compiled from images F^i-xii^ and G^i-x^. Melanin deposits (arrowheads).

To elucidate the cause of the female-specific melanization, we hypothesized that copulation may be involved, as males grasp females with their forelegs during mating using melanized bristles known as sex combs (Massey et al., 2019). First, to determine if melanization was copulation-dependent, virgin *epi>Hayan* female flies were collected. Additional control strains were used including a white-eyed mutant (*w*^*1118*^) commonly used as a *wild-type* control, *epi>, UAS-Hayan*, and *epi >Hayan*^*S322A*^ (a catalytically-inactive form of Hayan carrying Ala^322^ instead of Ser^322^ amino acid) (Bailey et al., 2020, Nam et al., 2012). None of the virgin female flies developed melanin, indicating that the melanization did not arise spontaneously (Figure 1D).

Next, we set up a nine-way mating scheme to determine whether the genetic background contributes to melanin formation. To do so, virgin female and male flies of the following strains were crossed: *epi>+* (heterozygous over *w*^*1118*^ strain; control), *epi>Hayan*, and *epi>Hayan*^*S322A*^. This created nine unique mating combinations, as each fly strain was mated with both females and males from the other two strains (Figure 1E). To link mating directly with melanization, one male and one virgin female fly were paired in a food vial, and copulation was visually confirmed. The successfully mated females were then separated and kept in isolation for further analysis. Interestingly, we found that female flies only developed melanin on their ventral abdomen if either partner in the mating pair expressed *Hayan* (Figure 1E). The melanin deposits were traced and superimposed onto a brightfield image of a virgin female abdomen to determine the size and pattern of melanization (Figure 1F and 1G). Melanin was more expansive when the female partner, rather than the male, expressed *Hayan* and appeared to localize to distinct regions on either side of the ventral midline (Figure 1E-1G and Figure S1A). Interestingly, melanin deposits consistently developed symmetrically adjacent to the 3^rd^ to 6^th^ tergites and varied in size from 89μm^2^ to 105,782μm^2^. Therefore, it appears that expression of *Hayan* in either the female or the male fly is sufficient to result in copulatory wounds to the female ventral abdomen.

Next, we sought to determine whether this melanization was specific to *Hayan* expression in the ventral fly epithelium or if it could be induced by producing the serine protease in other tissues. To do so, *Hayan* was expressed in the muscle (*MEF2-Gal4*), hemocytes (*Hml-Gal4*), fat body (*Cg-Gal4*), intestine (*esg-Gal4*), and with another epithelial driver (*NP2108-Gal4*) (Figure S1B, S1C, and Table S1). Virgin females for all fly strains were collected and melanization was not observed, except in the *NP2108>Hayan* strain (Figure S1B). The virgin *NP2108>Hayan* female flies developed spontaneous melanin deposits across their abdomens and had to be removed from further consideration as copulation-dependent melanization could not be distinguished. The remaining tissue-specific *Hayan* expressing strains were mated with their male counterparts (Figure S1B and S1C). Ultimately, at 3 days post copulation, only *epi>Hayan* flies developed melanin deposits. Thus tissue-specific expression of *Hayan* in other cell types did not result in copulatory wounds.

The GMR51F10-Gal4 (*epi>*) driver used here was obtained from the Janelia Gal4 collection, which was reported to drive expression in select neurons of the brain and ventral nerve cord (VNC) as well as in the adult fly ventral epithelium (Jenett et al., 2012). To determine whether Hayan activity in the *Drosophila* brain contributed to copulation-dependent melanization, *Hayan* was expressed with the pan-neural driver, *Elav-Gal4* (Table S1). As seen in Figure S1D, neuronal expression of *Hayan* was not sufficient to induce the melanin deposits. Next, we used *Elav-Gal80* to inhibit Gal4 transcriptional activity (Gal80 is an inhibitor of Gal4), which would prevent *Hayan* expression in neuronal cells, while preserving epithelial-specific expression (Figure S1E and Table S1). Melanization still occurred following mating with a strain harboring both *Elav-Gal80* and *epi>Hayan* as considerable melanin deposits were observed on female ventral abdomens post copulation (Figure S1E). We conclude that copulation-dependent melanization appears to be induced specifically by *Hayan* expression with *epi-Gal4* driver rather than in neurons or in the other cell types tested.

Next, we sought to determine how copulation or copulatory behavior could cause an injury-like response. Given the functional importance of sex combs on the male foreleg to mount and initiate mating, we first examined the male forelegs and observed that *epi>Hayan* male flies had noticeably longer bristles (Figure 2A and 2B) (Massey et al. 2019). The alteration in sex comb length suggests that the *epi>* driver is expressed in the epithelium of fly legs as well. Next, we assayed whether *Hayan* expression affected mating duration, however no significant difference was observed (Figure 2C). In visualizing the contact between the flies, we noticed that the male mid-legs made contact repeatedly, sometimes rapidly rubbing or poking the ventral abdomen in areas that correlated with the melanin deposits (Figure 2D, 2E, S1A, Movie S1 and S2). The cuticle and bristle pattern along the male mid-legs appeared indistinguishable between the *epi>+* and *epi>Hayan* flies and both mating pairs exhibited the mid-leg rubbing/poking behavior along the female ventral abdomen (Figure 2F, Movie S1 and S2). Nonetheless, we could not exclude the possibility of a subtle change that was not observable, such as a change in the rigidity or stiffness of the cuticle along the mid-legs that is the cause of injury to the female flies.

**Figure 2.**
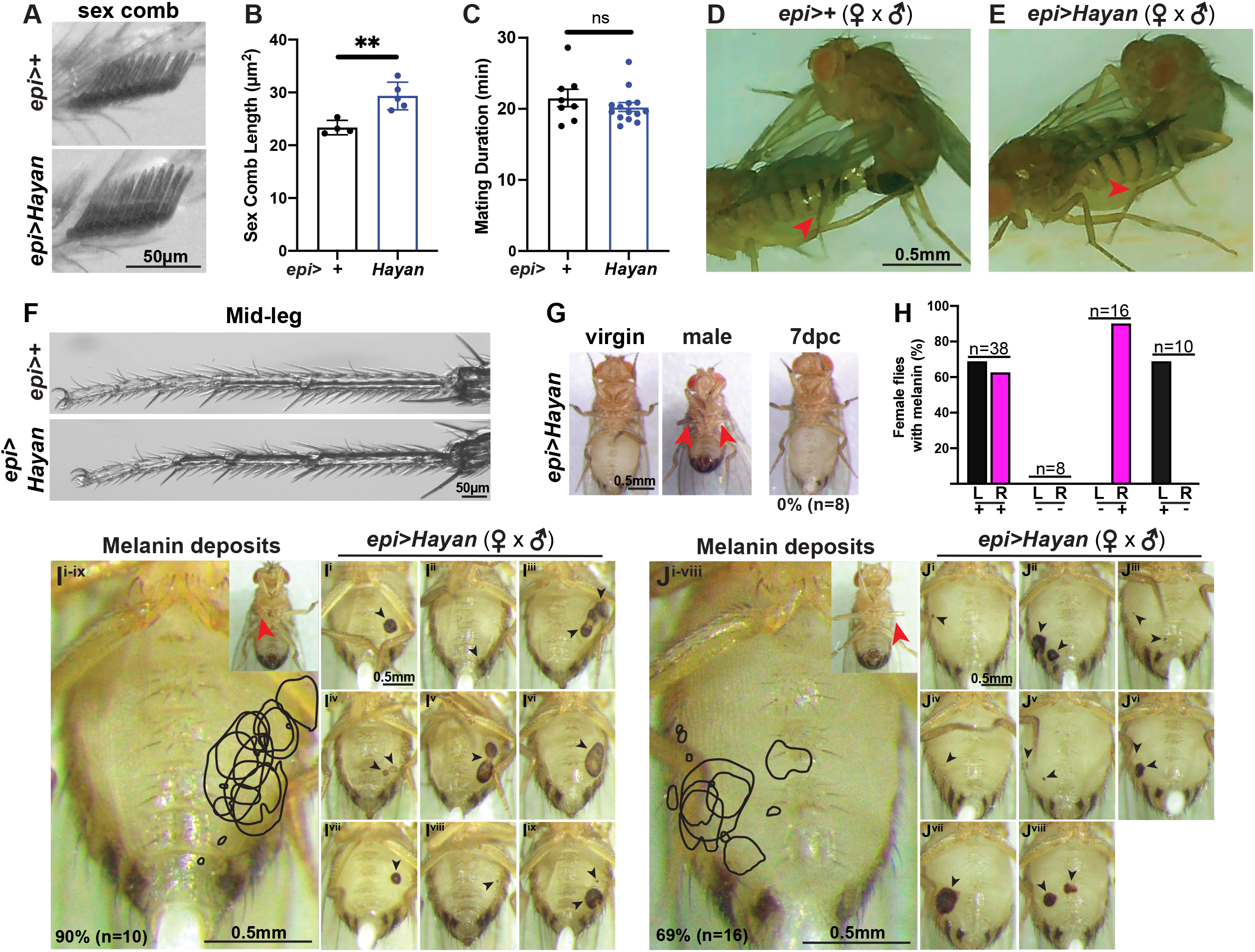
Copulation-induced melanization is dependent on male mid-leg contact with the female ventral abdomen. (A) Brightfield images of sex comb on male forelegs. (B) Quantification of sex comb length and (C) mating duration in strains indicated. Data represents the SEM with significance determined by Student T-test with P-value **<0.01. (D and E) Snapshot images of the male fly mid-leg contact with the female fly ventral abdomen (red arrowhead) during copulation. (F) Images of male fly mid-legs and (G) *epi>Hayan* mating pair with virgin female crossed to male with both mid-legs (red arrowheads) removed. No melanin arises on female abdomen post copulation with both mid-legs removed. (H) Quantification of percent melanized females post copulation. Denoted are the left (L) and right (R) sides of the female ventral abdomen and whether the male fly’s corresponding mid-leg was present (+) or removed (-). (I and J) Melanized regions of interest (ROI) from mating pairs indicated. The ROIs were compiled from images I^i-ix^ and J^i-viii^. Mid-leg removed (red arrowhead) and melanin deposits (black arrowheads).

Next, to directly assess the role of the male mid-legs in copulation-dependent melanization, we removed the mid-legs surgically. Mid-legless *epi>Hayan* males were allowed to mate with virgin *epi>Hayan* females, and copulation was confirmed by the presence of progeny in the food vials. However, none of the *epi>Hayan* female flies developed abdominal melanin (Figure 2G and 2H). To further confirm that male mid-legs were responsible for the melanin deposits, either the right or left mid-leg was surgically removed prior to copulation (Figure 2H-2J). Remarkably, the female *epi>Hayan* flies only developed melanin along the abdominal side that corresponded to the intact male mid-leg. Therefore, the expression of *Hayan* within the female ventral epithelium and/or male mid-legs appear to alter theses external structures such that they injure the female fruit fly during copulation.

In *Drosophila* and other insects, melanization is usually generated upon tissue damage at the site of injury (Galko and Krasnow, 2004, Losick et al., 2013, Ramet et al., 2002). Under laboratory conditions, melanin deposits can be induced following a needle puncture wound to the fly abdomen. These puncture wounds are reminiscent of copulatory wounds. As such, we were curious whether the mating induced melanin corresponded to an injury-like response. Our prior studies determined that the *Drosophila* epithelium repairs abdominal wounds through the generation of multinucleated, polyploid cells that arise via cell fusion and the endocycle, an incomplete cell cycle (Figure 3A) (Losick et al., 2013). To do so, *epi>Hayan* male and virgin females were allowed to mate for 1-2 days, after which the female flies were separated and dissected at 7 days post copulation. Large puncture wounds (3,000-10,000 μm^2^) can heal within 3 days post injury (dpi), but the copulation-induced melanin regions were up to ∼100,000 μm^2^ (Figure 3 and S2A). Therefore, additional time (7 days post copulation) was allowed to permit the copulatory wounds to heal.

**Figure 3.**
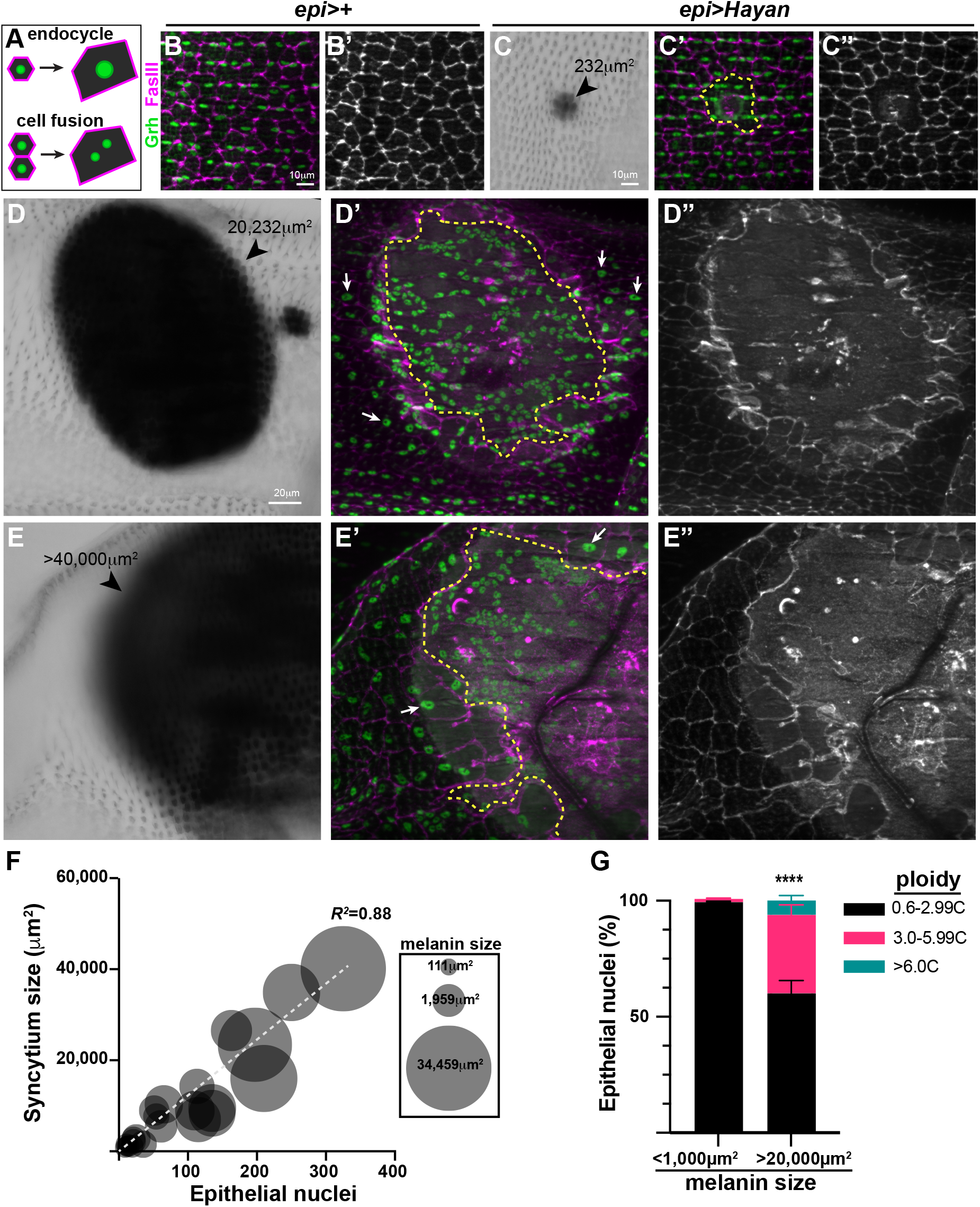
Multinucleated, polyploid cells arise at the site of melanization. (A) Illustration of the mechanisms of polyploidization via the endocycle (top) and cell fusion (bottom). (B-E) Representative immunofluorescent images of the epithelium and brightfield images of varying sizes of the melanin deposits generated in the mated *epi>Hayan* female flies. Melanin deposit (arrowhead), examples of enlarged epithelial nuclei (arrows), and the syncytium (yellow dashed line). Epithelial cell junctions (FasIII, magenta) and nuclei (Grh, green). (F) Quantification of the syncytium size as well as epithelial nuclear ploidy at 7dpc. Data represents the SEM with significance determined by Student T-test with P-value ****<0.0001.

The epithelium was visualized below the cuticle by immunostaining for two epithelial-specific proteins, FasIII, a septate junction protein, and Grh, a transcription factor that localizes to the epithelial nuclei (Figure 3B). The epithelium in the *wild-type* strains, including the *epi>+* used in this study, is made up of mononucleated, diploid cells (Losick et al., 2013). We observed a central multinucleated cell known as a syncytium in epithelium underlying the melanin deposits following copulation. The size of the syncytium correlated with the size of the melanized area (Figure 3C-3E). As expected, when syncytium size increased, so did the number of epithelial nuclei (Figure 3F, *R*^*2*^=0.88) (Losick et al., 2013). Copulation generated melanin areas varied dramatically in size with the majority either being ≤1,000 μm^2^ or ≥20,000μm^2^. For small melanin areas, we noticed the underlying nuclei remained unchanged in size, whereas the epithelial nuclei within and surrounding the syncytium from large melanin deposits were enlarged. Epithelial nuclear DNA content was measured to verify that the increase in nuclear area was indicative of a change in nuclear ploidy (see Methods). Indeed, nuclei surrounding the large melanin (>20,000μm^2^) areas had a significant increase in nuclear DNA content, which ranged from 3C to 12C, whereas nuclei below smaller melanin deposits remained diploid (2C) and did not endocycle (Figure 3G).

To investigate whether there is a size threshold required to increase nuclear ploidy, we utilized the needle puncture wound method (Bailey et al. 2020). Similar to copulatory wounds, we found that the epithelial nuclei surrounding small puncture wounds with melanin areas ≤1,000μm^2^ remained diploid (Figure S2). In comparison, areas that reached ≥3,000 μm^2^ became polyploid with epithelial nuclear ploidy ranging from 3C to 12C. Melanin regions of all sizes generated by a needle puncture wound formed a central syncytium at the site of injury. Therefore, cell fusion is initiated for any sized injury, whereas endoreplication requires a threshold of epithelial nuclei loss. Interestingly, we also observed a similar response in a genetic-induced damage model through conditional expression of the apoptotic gene, *hid*, in the fly epithelium (Shen et al., 2026). Enhanced cell death was necessary to induce endoreplication in this genetic model of tissue injury. Thus, the *Drosophila* epithelium utilizes both cell fusion and the endocycle to heal a variety of insults. Moreover, the generation of multinucleated cells through cell fusion appears to function as the tissue’s initial defense against damage. When an injury exceeds a compensatory threshold, the tissue then utilizes the endocycle to sufficiently respond to greater damage and compensate for cell loss.

In sum, we have shown that overexpression of the serine protease, *Hayan*, renders *Drosophila melanogaster* susceptible to injury during copulation. This wounding is dependent on contact with male fly mid-legs and healed through the generation of polyploid cells. Cell fusion is initiated for small wounds, but the endocycle is necessary to compensate for more substantial cell loss. Therefore, we speculate that polyploidization evolved as a mechanism to efficiently repair tissue damage caused by copulatory wounds in nature.

## Data Availability Statement

The authors affirm that all data necessary for verifying the conclusions are present within the article, figures, and Supplemental information. All raw data points reported in figures can be found in the Source data file. Additionally, the *Drosophila* strains and antibodies used in this study are listed Supplementary Table 1 and 2 and are available upon request or as referenced.

## Acknowledgements

We would like to thank the members of the Losick Lab, including Stefania Bonanni, Minqi Shen, and Jack Weidenbach for critical review of this manuscript. We also thank Madisen Caferro who first observed that overexpression of *Hayan* caused melanization and Daria Perminova who observed that melanization is dependent on copulation during their graduate rotation projects in the Losick Lab. Walker Vickers (Das Lab at BC) assisted with video editing and Bret Judson as well as the Boston College Imaging Core for infrastructure and support. Finally, a special thanks to Dr. Michael Crickmore (Harvard Medical School) who provided use of his imaging equipment and mating chambers. The fly strains used in this study were purchased from (with support by): the Bloomington Drosophila Stock Center (NIH P40OD018537). The FasIII antibody used in this study was purchased from the Developmental Studies Hybridoma Bank (created by the NICHD and maintained at The University of Iowa, Department of Biology, Iowa City, IA 52242, USA).

## Study Funding

Research reported in this publication was supported by Boston College and the National Institute of General Medical Sciences of the National Institutes of Health under Award Number R35GM124691 to V.P.L. The content is solely the responsibility of the authors and does not necessarily represent the official views of the National Institutes of Health.

## Competing Interest Statement

The authors declare no competing or financial interests.

## Author Contributions

Conceptualization: S.L.J., L.M.B, and V.P.L.; Methodology: S.L.J., L.M.B, M.J.P, and V.P.L.; Validation: S.L.J., L.M.B, and V.P.L.; Formal analysis: S.L.J., L.M.B, and V.P.L.; Investigation: S.L.J., L.M.B, M.J.P., and V.P.L.; Resources: M.J.P. and V.P.L. Writing - original draft: S.L.J. and V.P.L.; Writing - review & editing: S.L.J., L.M.B, M.J.P., and V.P.L.; Visualization: S.L.J., L.M.B, and V.P.L.; Supervision: V.P.L.; Project administration: V.P.L.; Funding acquisition: V.P.L.

